# A Longitudinal Model for Functional Connectivity Networks Using Resting-State fMRI

**DOI:** 10.1101/152538

**Authors:** Brian Hart, Ivor Cribben, Mark Fiecas, for the Alzheimer’s Disease Neuroimaging Initiative

## Abstract

Many neuroimaging studies collect functional magnetic resonance imaging (fMRI) data in a longitudinal manner. However, the current network modeling literature lacks a general framework for analyzing functional connectivity (FC) networks in fMRI data obtained from a longitudinal study. In this work, we build a novel longitudinal FC network model using a variance components approach. First, for all subjects’ visits, we account for the autocorrelation inherent in the fMRI time series data using a non-parametric technique. Second, we use a generalized least squares approach to estimate 1) the within-subject variance component shared across the population, 2) the FC network, and 3) the FC network’s longitudinal trend. Our novel method for longitudinal FC networks seeks to account for the within-subject dependence across multiple visits, the variability due to the subjects being sampled from a population, and the autocorrelation present in fMRI data, while restricting the number of parameters in order to make the method computationally feasible and stable. We develop a permutation testing procedure to draw valid inference on group differences in baseline FC and change in FC over time between a set of patients and a comparable set of controls. To examine performance, we run a series of simulations and apply the model to longitudinal fMRI data collected from the Alzheimer’s Disease Neuroimaging Initiative database.

## 1 Introduction

### 1.1 Resting-State fMRI

Resting-state functional magnetic resonance imaging (fMRI) captures a series of images of the brain in subjects who are not given a particular task to perform while in the scanner. The scanner repeatedly captures blood oxygenation level dependent (BOLD) signals at hundreds of thousands of locations within the brain, creating a time series of images of the brain. By capturing the BOLD signal of the resting brain, resting-state fMRI provides an opportunity for researchers to examine the functional connectivity (FC) network not tied to a particular task. We define FC as the temporal dependence, measured through cross-correlations, in the BOLD signals between brain regions [Friston et al., 1993]. Identifying group differences in FC can help better understand the underlying neurological process of a disease and its progression. Observed group differences can also potentially form biomarkers to be used for early detection and treatment of neurological disorders [Fox and Raichle, 2007].

### 1.2 Alzheimer’s Disease Neuroimaging Initiative Data

A subset of the Alzheimer’s Disease Neuroimaging Initiative (ADNI) data was used to demonstrate a practical application of our model. The data consists of longitudinal resting-state fMRI images collected at baseline, 3 months from baseline, 6 months from baseline, 12 months from baseline, and annually thereafter. There are two groups of interest, the cognitively normal (CN) group and the Alzheimer’s Disease (AD) group. We focused our attention on late-onset AD and included only patients who were 65 years of age or older at baseline [van der Flier et al., 2011, Holland et al., 2012]. To better separate the AD and CN groups, only patients who remained in one group for the entirety of the follow-up were considered in our analysis. The remaining CN group consists of 111 visits from 30 patients (17 females and 13 males) with each patient having between 1 and 6 visits. The AD group consists of 79 visits from 26 patients (11 females and 15 males) with each patient having between 1 and 5 visits. The average age was 75.9 for the CN group with a range of 65.2 to 95.7, while the AD group average age was 76.7 with a range of 66.5 to 88.6.

**Figure 1:**
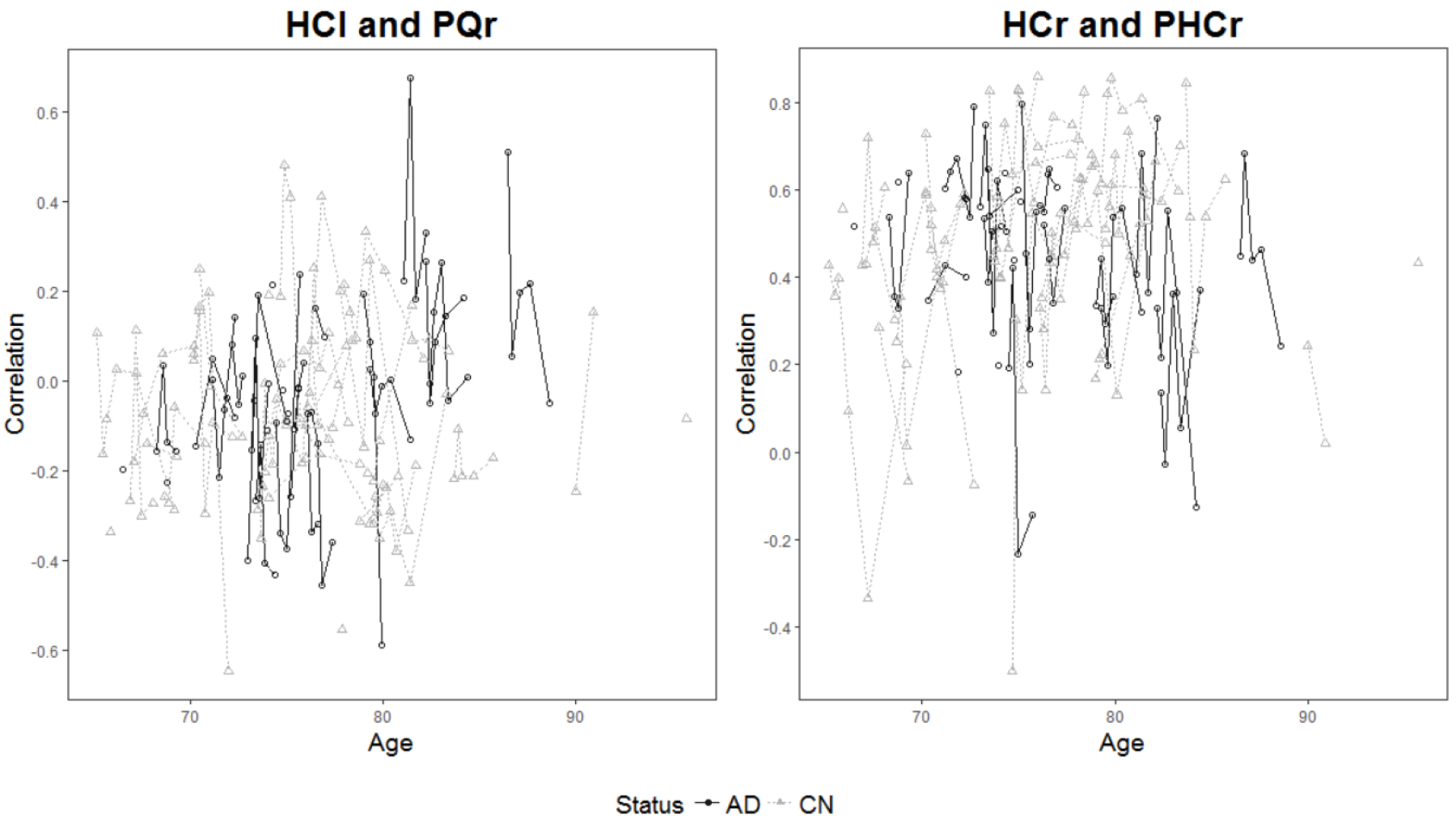
Spaghetti plots of the correlation between two ROIs against age. Each point represents a visit, and each line represents a subject. The ROIs represented in these plots are the left and right hippocampus (HCl and HCr), right precuneus (PQr), and right parahippocampus (PHCr).

### 1.3 Clinical Motivations

Previous works have demonstrated the utility of FC network analysis. For example, past research has identified altered FC between healthy aging patients and those with AD. Even among CN individuals, FC demonstrates aging effects that are heterogeneous between different brain regions [Chen et al., 2016]. Chase [2014] and Hafkemeijer et al. [2012] showed altered FC patterns beyond healthy aging in patients with dementia and AD. Other works, including Ren et al. [2016] and Wang et al. [2007] have noted abnormal FC in various stages of AD. Wang et al. [2012] even demonstrated the impact of family history of AD on FC. In addition, Xiang et al. [2013] showed decreased FC from CN patients to mild cognitively impaired (MCI) patients to AD patients, and Li et al. [2015] found decreased network FC for CN patients who progressed to MCI over the following 24 months compared to non-progressers. These previous works, however, used *cross-sectional models*. Our work will build off of these results on the clinical utility of FC as a biomarker for AD.

### 1.4 Methodological Considerations

Of the studies mentioned in Section 1.3, only Ren et al. [2016] utilized truly longitudinal fMRI data. Aging effects are often measured by comparing young and ederly groups rather than following one group of subjects over time. A comprehensive longitudinal model that tests for differences in baseline and trend is needed to verify and expand on the previous results. Finn and Constable [2016] demonstrated that CN patients have distinct brain signatures in fMRI images, implying that separate scans from a single individual exhibit dependence. This finding can be leveraged in a longitudinal framework to better model aging effects. Our new methodology analyzes fMRI data at the region of interest (ROI) level. Thus, before fitting the model, the investigator must select a number of ROIs to include in the FC network analysis. We motivate the challenges of longitudinal FC network analysis with a preliminary examination of the ADNI data. Figure 1 shows spaghetti plots of the FC between the preprocessed fMRI time series obtained from two ROI pairs for the AD and CN groups. The clustering of points within each line shows the within-subject dependence. In addition, there is considerable within-subject and within-group noise present in the estimates of FC. What is not evident from the figure is that the time series from which these correlations were obtained exhibit autocorrelation that contributes to the overall variability in FC. To add another level of complication, the figure depicts the marginal relationship between two ROIs, but to properly model the entire network we need a joint model that considers the network of all possible pairwise groupings of the chosen ROIs.

While methods such as Fiecas et al. [2017] exist for the cross-sectional case, no modeling framework exists for longitudinal FC network analysis. We fill this gap in the literature by proposing a novel longitudinal model and inference procedure that considers the network of all possible pairwise groupings of the chosen ROIs in resting-state fMRI data. Our longitudinal variance components FC network model accounts for the within-subject dependence across multiple visits, the variability due to the subjects being sampled from a population, and any autocorrelation present in fMRI time series. We also propose an efficient permutation based inference procedure that allows for valid hypothesis testing of group differences in baseline FC networks and FC network aging effects.

The remainder of the paper is laid out as follows. Section 2 formally introduces the model, including the estimation and inference procedures. Section 3 explains the design of the simulation study and discusses the results. Section 4 more fully describes the ADNI data, specifies the models fit to the data, and interprets the results of the FC analysis. We close with a discussion of future work and a conclusion in Section 5. R code for the methods proposed in this paper may be found at https://github.com/mfiecas/longitudinalFC.

## 2 Materials and Methods

### 2.1 Model Specification

Suppose we have a cohort of *N* individuals and let *P* denote the number of ROIs selected for a FC network analysis. We collect a *P*-variate, fMRI time series of length *T* from the preprocessed fMRI images of each of the *N* subjects at each visit. Let the subscripts *i* and *j* denote subject and visit, respectively. A particular subject returns for *J*_*i*_ total visits, and the cohort has a total of 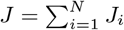 visits. Let *Y*_*i*_ represent the vector of sample correlation coefficients for subject *i* of length *QJ*_*i*_, where *Q* = *P*(*P* – 1)/2 is the number of ROI pairs. Within *Y*_*i*_, the *Q* correlations from the first visit, *Y*_*i*1_, are followed by the *Q* correlations from the second visit, *Y*_*i*2_, and so on until the *J*_*i*_-th visit. The full response vector *Y* is formed by stacking the *N* different *Y*_*i*_ vectors. Our longitudinal model for the FC network is a linear model with baseline effect *β*_0_ and longitudinal trend *β*_1_, where each of these model parameters are vectors of length *Q*. We denote the time at visit *j* for subject *i* as *v*_*ij*_. The vector *v*_*i*_ is formed by stacking the *J*_*i*_ distinct 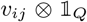 vectors for subject *i*, where 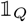 is a vector of ones of length *Q*. Likewise *v* is formed by stacking the *N* distinct *v*_*i*_ vectors. Depending on the nature of the data and the research questions at hand, *v*_*ij*_ can be set to the visit number, the time since baseline, or the patient’s age. Then, denoting element-wise multiplication with ∗, our model has the following linear form:

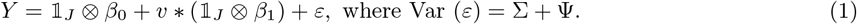

The key element in our longitudinal linear model is the variance structure of the error term. We separate the error variance into two components, Σ and Ψ. Σ accounts for the within-visit variance and the temporal autocorrelation in the fMRI time series, and Ψ accounts for the variability and covariability arising from the heterogeneity across subjects and the within-subject covariation coming from the longitudinal design. Σ is block diagonal where each *Q* × *Q* block, Σ_*ij*_, accounts for the within-visit variance present in visit *j* for subject *i* for the *Q* pairs of ROIs. Ψ is also block diagonal with a *QJ*_*i*_ × *QJ*_*i*_ block for participant *i*. These diagonal blocks do not differ between subjects except through their dimensions which depends on the number of visits for each subject. Let Ψ_*diag*_ be an arbitrary diagonal block of Ψ. We then further break Ψ_*diag*_ into two components, Ψ_0_ and Ψ_1_. Ψ_0_ is a *Q* × *Q* block that is repeated along the diagonal of each Ψ_*diag*_. This term models the within-visit covariability not captured by Σ. Ψ_1_ is a *Q* × *Q* block that populates the off diagonal blocks of Ψ_*diag*_, modeling the within-subject, across-visit covariability coming from the longitudinal design.

We write Equation 1 in the form of a linear model with a vector response, allowing us to use existing methods for estimating the parameters and for statistical inference. To this end, our model can also be written in the standard linear model form with a design matrix *X*. Let *X*_*ij*_ = [1 *v*_*ij*_] ⊗ *I*_*Q*_, where *I*_*Q*_ is the *Q* × *Q* identity matrix. To form *X*_*i*_, the portion of the design matrix specific to subject *i*, we stack the *J*_*i*_ individual *X*_*ij*_. Likewise, to form *X* we stack the *N* individual *X*_*i*_. If we define *β* as a vector of length 2*Q* where the first *Q* elements are *β*_0_ and the last *Q* elements are *β*_1_, then Equation 1 can be written as *Y* = *Xβ* + *ε*.

Figure 2 shows a workflow chart of the procedure used to estimate the model parameters and test hypotheses which are subsequently described in more detail.

**Figure 2:**
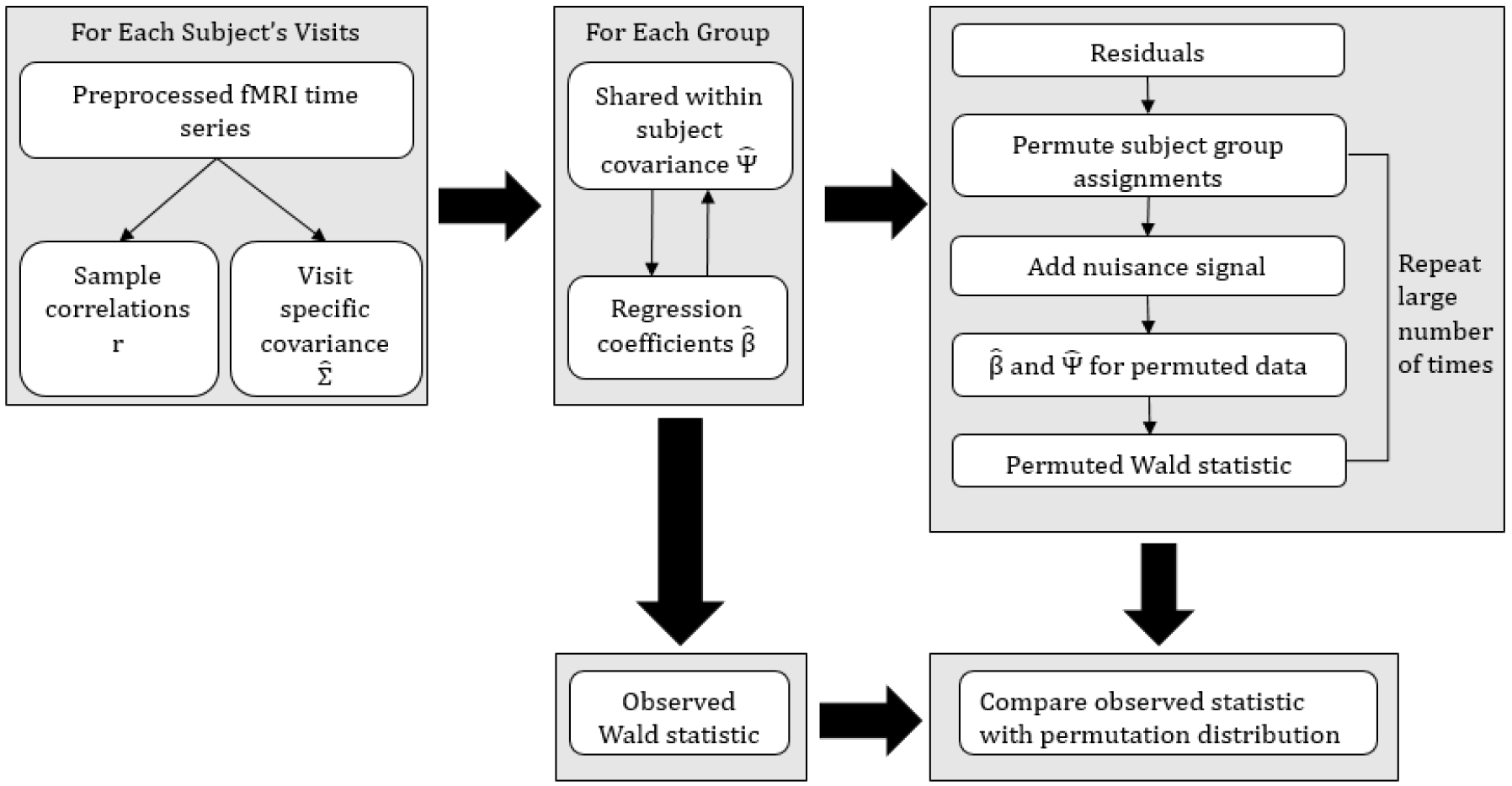
A workflow chart of the estimation and inferential procedure of our variance components model.

### 2.2 Estimating Within Visit Covariance

We start by estimating the sample correlation coefficient between all ROI pairs for all visits and their corresponding variances and covariances. Let (*w*_1*t*_,…, *w*_*pt*_)′ for *t* ∈ {1, 2,…,*T*} be the time series of preprocessed BOLD signals from *P* ROIs for a single visit. Here the first subscript on *w* indicates the ROI and the second subscript indicates the time within the time series. Then for the *p*-th and *q*-th ROIs, where *p*, *q* ∈ {1, 2,…, *P*}, we have 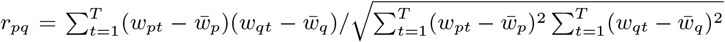. We now seek a consistent estimator of the covariance between correlations of two ROI pairs, (*p,q*) and (*p*′,*q*′). The estimator of Melard et al. [1991] and Roy [1989] offers a nice solution which accounts for the autocorrelation present in fMRI time series and is consistent under mild conditions. First, define 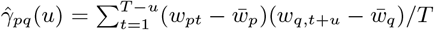. Then, letting *h*(·) be the modified Bartlett window with bandwidth *b*(*T*), we set 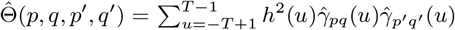. Using 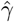 and 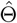, we then let 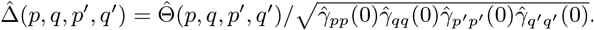. Finally, our estimator takes the following form:

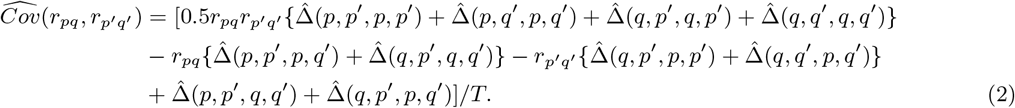

We use Equation 2 to populate the cells of the each Σ_*ij*_ block to get our estimates 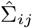.

### 2.3 Estimating Between Subject Covariance, Ψ, and *β*

Utilizing a generalized least squares (GLS) approach, we now proceed with the estimation of the between subject covariance Ψ and the regression coefficients *β*, conditional on the previously estimated within-visit covariances, 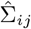.

Although the framework allows for many different structures for Ψ, we assume a block compound symmetry structure so that all diagonal blocks, Ψ_0_, are equal and all off diagonal blocks, Ψ_1_, are equal. The block compound symmetry assumption keeps the parameter space to a reasonable size, but note that one could easily consider other forms of Ψ, such as an autoregressive structure, with minimal modification to the estimation procedure. We use the ordinary least squares estimator 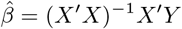 to provide a good starting estimate of *β*. We then update the two components of Ψ using the following formulas:

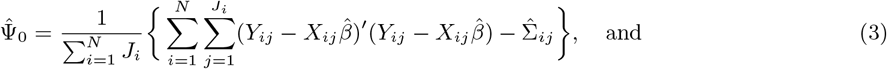

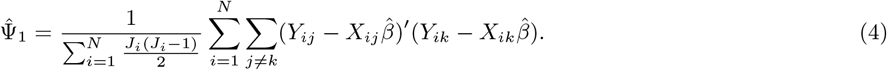

Note how these estimators resemble the mean squared error. For 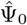, the sum of the squared residuals for all visits are summed. We then subtract the previously estimated 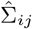 and divide the remaining covariance matrix by the total number of visits to get 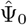. We estimate Ψ_1_ in a similar fashion. In this case, we calculate and average the cross products of the residuals from all pairs of visits for each subject. We do not subtract a 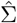 term here because 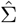 has been set to 0 for the off diagonal blocks occupied by Ψ_1_. These simple estimators, obtained through matching moments, significantly reduce computation in comparison to a maximum likelihood approach. To increase model parsimony, different structures can be considered for Σ_*ij*_, Ψ_0_, and Ψ_1_. For example, to enforce a diagonal structure, set all off-diagonal elements to 0, or to enforce a compound symmetry structure, set all diagonal elements to the average of the diagonal elements and likewise for the off-diagonal elements [Laird, 2004]. Changing the form of Σ_*ij*_, Ψ_0_, and Ψ_1_ allows you to fit the model with flexible variance assumptions as is often done in traditional generalized least squares linear models.

With an estimate of Ψ obtained using Equations 3 and 4, we can now use the standard GLS formula to update the regression coefficients as follows: 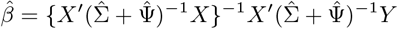. At this point we have two choices: iteratively update 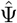 and 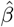 until convergence (full convergence), or accept the estimates (one-step) and proceed with the inferential procedure. The one-step estimator provides a significant advantage in computing time as 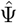 and 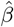 must be estimated for each permutation of the inference procedure [Ganjgahi et al., 2015]. Additionally, Amemiya [1977] proved that the one-step GLS estimator maintains consistency.

We estimate 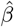, 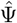, and 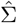 for each group (CN and AD for ADNI) separately using this estimation procedure. Superscripts on the parameter estimates denote the group (e.g. 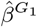, 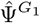, and 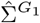 are the estimates for group 1).

### 2.4 Inference

Two general hypothesis tests are of interest in a longitudinal FC model: the group difference in the baseline FC, and the group difference in the longitudinal trend in FC. For each hypothesis, we would like to test for the significance of the group difference in both the global FC network and the local ROI pair FC. We refer to the vector wide test of a difference in the parameter vector *β*_0_ or *β*_1_ as a global test and refer to a test of a group difference in a single element of *β*_0_ or *β*_1_ as a local test. To accomplish our hypothesis testing objectives, we use the Wald statistic, 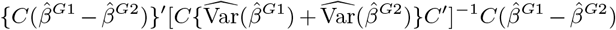, and adjust the contrast matrix, *C*, depending on the hypothesis of interest. For instance, to test for a global difference in *β*_0_, we replace all *β* terms with the *β*_0_ vector for the proper group and set the contrast matrix, *C*, to the *Q* × *Q* identity matrix.

We estimate the variance of each group’s regression coefficients using 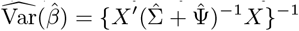. We now detail a permutation procedure for all inference.

1. Calculate residuals from the fitted model for each subject: 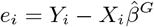 for subject *i* in group *G*.
2. Permute group assignments of *e*_*i*_.
3. Add the nuisance signal back to *e*_*i*_ based on new permuted group assignments *G*\*. For the main effect (intercept) tests we add in the longitudinal trends by setting 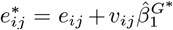. Likewise, for the interaction (slope) tests we set 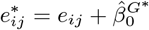.
4. Refit the model on *e*^*^, the permuted, adjusted, and stacked residuals from step 3.
5. Calculate a new Wald statistic for the fitted values of 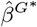 and 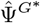.

We repeat steps 2 through 5 a large number of times to create a permutation distribution to be used as a reference distribution of the originally calculated test statistic. The number of permutations determines the precision of the *p*-value and should be chosen to be large enough to offer sufficient precision after any multiple comparisons adjustment. In our particular data example we chose to run 5,000 permutations. Because the obtained *p*-values are estimated discrete values, we additionally use the permutation *p*-value correction procedure of Phipson et al. [2010]. To account for the fact that 2*Q* local hypotheses are tested simultaneously, we then apply the false discovery rate (FDR) controlling procedure of Benjamini and Hochberg [1995] to the corrected *p*-values from the local tests. The Phipson et al. [2010] correction helps avoid unadjusted *p*-values with value 0 which may improperly maintain significance after a multiple comparisons correction.

Chung and Romano [2013] showed that studentized test statistics, such as the proposed Wald statistic, allow for valid inference in many permutation test settings. A recent comparison of the performance of different permutation strategies by Winkler et al. [2014] showed that the Ter Braak permutation testing procedure we use maintains nominal Type I error and is fairly robust [Ter Braak, 1992]. This method offers the additional advantage that the data only needs to be permuted once and the model only fit twice at each iteration of the permutation test to test all local and global hypotheses. Testing all hypotheses under a single permutation schedule greatly reduces the computational burden of the testing procedure.

## 3 Simulation Study

### 3.1 Simulation Setup

A series of simulations were designed with different data generating mechanisms to assess model performance. In all scenarios each time series contained 120 time points and had an autocorrelation structure that followed a first-order autoregressive process with an AR parameter of 0.3. A multivariate time series was simulated for each subject at three visits. For each visit, the Q correlations were simulated from a multivariate normal distribution where the mean and variance varied by group based on the simulation setting. For group 1, the mean vector was always assumed to be 0 and the covariance matrix was the same across all simulation settings. The simulations used *P* of either 3, 5, or 10 as the dimension of the multivariate normal distribution. For the 3 and 5 dimension settings only the first element of the group 2 mean vector was allowed to vary by simulation setting, while the other elements were set to match group 1. For the 10 dimension settings the first 5 elements of the group 2 mean vector varied by simulation setting and the other elements were again set to match group 1. 1,000 Monte Carlo simulations were run for all simulation settings with 3 and 5 dimensions, and 500 simulations were run for the 10 dimensional simulation settings. Group sizes of 15 and 30 were considered. The true variance of the correlations was either equal for the two groups or the group 2 variance was double the group 1 variance. 500 permutations were used for the permutation test for all settings. The group size, number of visits, and time series length were selected to reflect values found in typical fMRI studies. The effect size of 0.1 is realistic for fMRI longitudinal scenarios. Considering a smaller effect size would lead to very similar conclusions with slight decreases in power across all models considered.

As mentioned earlier, there are no other competitor models which consider FC networks longitudinally. Because of this, we have chosen to fit three versions of our model with different variance assumptions and estimation methods so they can be compared to each other. The first model considered was a full convergence model which iterated between 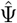 and 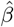 until convergence. It assumed an unstructured Σ_*ij*_ and compound symmetry for Ψ_0_ and Ψ_1_. This model is referred to as the full convergence full variance model. The second was a one-step model which stops after one iteration of solving for 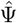 and 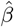. It also assumed an unstructured Σ_*ij*_ and compound symmetry for Σ_0_ and Σ_1_ and is referred to as the one-step convergence full variance model. The final model was a one-step model which assumed a diagonal structure for Σ_*ij*_ and scaled identity structures for Σ_0_ and Σ_1_. This last model is referred to as the one-step convergence reduced variance model.

### 3.2 Simulation Results

Tables 1 and 2 show the simulation study results. Table 1 shows the global and local Type I error for the main effect and interaction across all simulations. The reported global test results are the average global Type I errors across 500 or 1,000 Monte Carlo runs. The local test results are the average Type I errors of the unadjusted *p*-values for all null hypotheses across the 500 or 1,000 Monte Carlo runs. While the local *p*-values would be adjusted in practice, the numbers in the table provide easy reference to a nominal Type I error of 0.05. Table 2 shows the average global power and average local power using false discovery rate adjusted *p*-values. All *p*-values were corrected using the method of Phipson et al. [2010]. Additional simulation study results can be found in Web Table 1 of web-based supplementary materials. Table 1 shows roughly nominal Type I error rates for all three model specification. While there was some slight inflation in all three models, especially for the 10 ROI simulations, the inflation was attenuated by the increase in sample size from 15 to 30 per group. Table 1 demonstrates adequate power, both locally and globally for all three models. As expected, power increased with larger group size and decreased with a larger true group 2 variance.

There was no consistent difference in performance between the three models in terms of power or Type I error across the different simulation settings. The two full variance settings match the true model of the simulated data, yet the reduced variance model did not suffer in comparison. The reduced variance model may have offered similar performance because the smaller parameter space allowed for improved estimation. The reduced variance model did not capture the full true variance, but it still performed well by allowing the FC for each ROI pair to be correlated across multiple visits for a given subject.

**Table 1:** Type I error rates for simulation study for all globally null simulation settings. Type I errors for the main effect (group difference in intercepts) and interaction effect (group difference in longitudinal slopes) are reported both globally and locally. The global Type I errors are averaged across all simulations. The local Type I errors reported are unadjusted and averaged across all simulations and all null ROI pairs.

|  |  | Convergence Type:<br>Variance Structure: |  |  | Full<br>Full |  | One-Step<br>Full |  |  | One-Step<br>Reduced |  |
| --- | --- | --- | --- | --- | --- | --- | --- | --- | --- | --- | --- |
| | Group<br>Size | Variance | $\beta_0$ | $\beta_1$ | 3<br>ROIs | 5<br>ROIs | 3<br>ROIs | 5<br>ROIs | 10<br>ROIs | 3<br>ROIs | 5<br>ROIs |
| Main<br>Effect<br>Global<br>Test | 15 | Equal | 0 | 0 | 0.061 | 0.052 | 0.057 | 0.055 | 0.066 | 0.056 | 0.054 |
|  |  |  | 0 | 0.1 | 0.065 | 0.065 | 0.067 | 0.071 | 0.064 | 0.067 | 0.063 |
|  |  | Group 2<br>Double | 0 | 0 | 0.058 | 0.049 | 0.055 | 0.053 | 0.054 | 0.056 | 0.048 |
|  |  |  | 0 | 0.1 | 0.068 | 0.058 | 0.069 | 0.056 | 0.068 | 0.070 | 0.060 |
|  | 30 | Equal | 0 | 0 | 0.045 | 0.049 | 0.046 | 0.050 | 0.066 | 0.043 | 0.047 |
|  |  |  | 0 | 0.1 | 0.046 | 0.050 | 0.054 | 0.047 | 0.066 | 0.048 | 0.049 |
|  |  | Group 2<br>Double | 0 | 0 | 0.057 | 0.054 | 0.058 | 0.053 | 0.062 | 0.058 | 0.060 |
|  |  |  | 0 | 0.1 | 0.049 | 0.066 | 0.055 | 0.059 | 0.078 | 0.058 | 0.053 |
| Main<br>Effect<br>Local<br>Test | 15 | Equal | 0 | 0 | 0.071 | 0.061 | 0.073 | 0.061 | 0.066 | 0.066 | 0.062 |
|  |  |  | 0 | 0.1 | 0.064 | 0.067 | 0.067 | 0.069 | 0.069 | 0.064 | 0.067 |
|  |  | Group 2<br>Double | 0 | 0 | 0.064 | 0.063 | 0.061 | 0.062 | 0.071 | 0.064 | 0.064 |
|  |  |  | 0 | 0.1 | 0.068 | 0.062 | 0.069 | 0.063 | 0.069 | 0.065 | 0.063 |
|  | 30 | Equal | 0 | 0 | 0.050 | 0.059 | 0.048 | 0.058 | 0.058 | 0.048 | 0.057 |
|  |  |  | 0 | 0.1 | 0.053 | 0.058 | 0.051 | 0.058 | 0.056 | 0.052 | 0.057 |
|  |  | Group 2<br>Double | 0 | 0 | 0.063 | 0.055 | 0.062 | 0.055 | 0.057 | 0.060 | 0.056 |
|  |  |  | 0 | 0.1 | 0.054 | 0.059 | 0.055 | 0.058 | 0.059 | 0.053 | 0.057 |
| Interaction<br>Global<br>Test | 15 | Equal | 0 | 0 | 0.062 | 0.054 | 0.052 | 0.057 | 0.072 | 0.049 | 0.052 |
|  |  |  | 0.1 | 0 | 0.046 | 0.054 | 0.044 | 0.049 | 0.076 | 0.044 | 0.049 |
|  |  | Group 2<br>Double | 0 | 0 | 0.057 | 0.067 | 0.050 | 0.062 | 0.070 | 0.051 | 0.069 |
|  |  |  | 0.1 | 0 | 0.079 | 0.071 | 0.078 | 0.061 | 0.090 | 0.073 | 0.063 |
|  | 30 | Equal | 0 | 0 | 0.054 | 0.059 | 0.058 | 0.061 | 0.078 | 0.056 | 0.058 |
|  |  |  | 0.1 | 0 | 0.052 | 0.067 | 0.049 | 0.063 | 0.070 | 0.051 | 0.051 |
|  |  | Group 2<br>Double | 0 | 0 | 0.068 | 0.065 | 0.071 | 0.061 | 0.072 | 0.075 | 0.057 |
|  |  |  | 0.1 | 0 | 0.056 | 0.041 | 0.057 | 0.049 | 0.068 | 0.059 | 0.040 |
| Interaction<br>Local<br>Test | 15 | Equal | 0 | 0 | 0.059 | 0.062 | 0.058 | 0.062 | 0.067 | 0.059 | 0.064 |
|  |  |  | 0.1 | 0 | 0.055 | 0.060 | 0.054 | 0.057 | 0.071 | 0.057 | 0.060 |
|  |  | Group 2<br>Double | 0 | 0 | 0.063 | 0.065 | 0.065 | 0.061 | 0.072 | 0.064 | 0.063 |
|  |  |  | 0.1 | 0 | 0.074 | 0.067 | 0.071 | 0.061 | 0.067 | 0.074 | 0.064 |
|  | 30 | Equal | 0 | 0 | 0.050 | 0.055 | 0.049 | 0.054 | 0.059 | 0.049 | 0.055 |
|  |  |  | 0.1 | 0 | 0.054 | 0.059 | 0.053 | 0.058 | 0.056 | 0.058 | 0.059 |
|  |  | Group 2<br>Double | 0 | 0 | 0.061 | 0.059 | 0.062 | 0.057 | 0.060 | 0.063 | 0.060 |
|  |  |  | 0.1 | 0 | 0.057 | 0.055 | 0.057 | 0.055 | 0.058 | 0.058 | 0.055 |

**Table 2:**
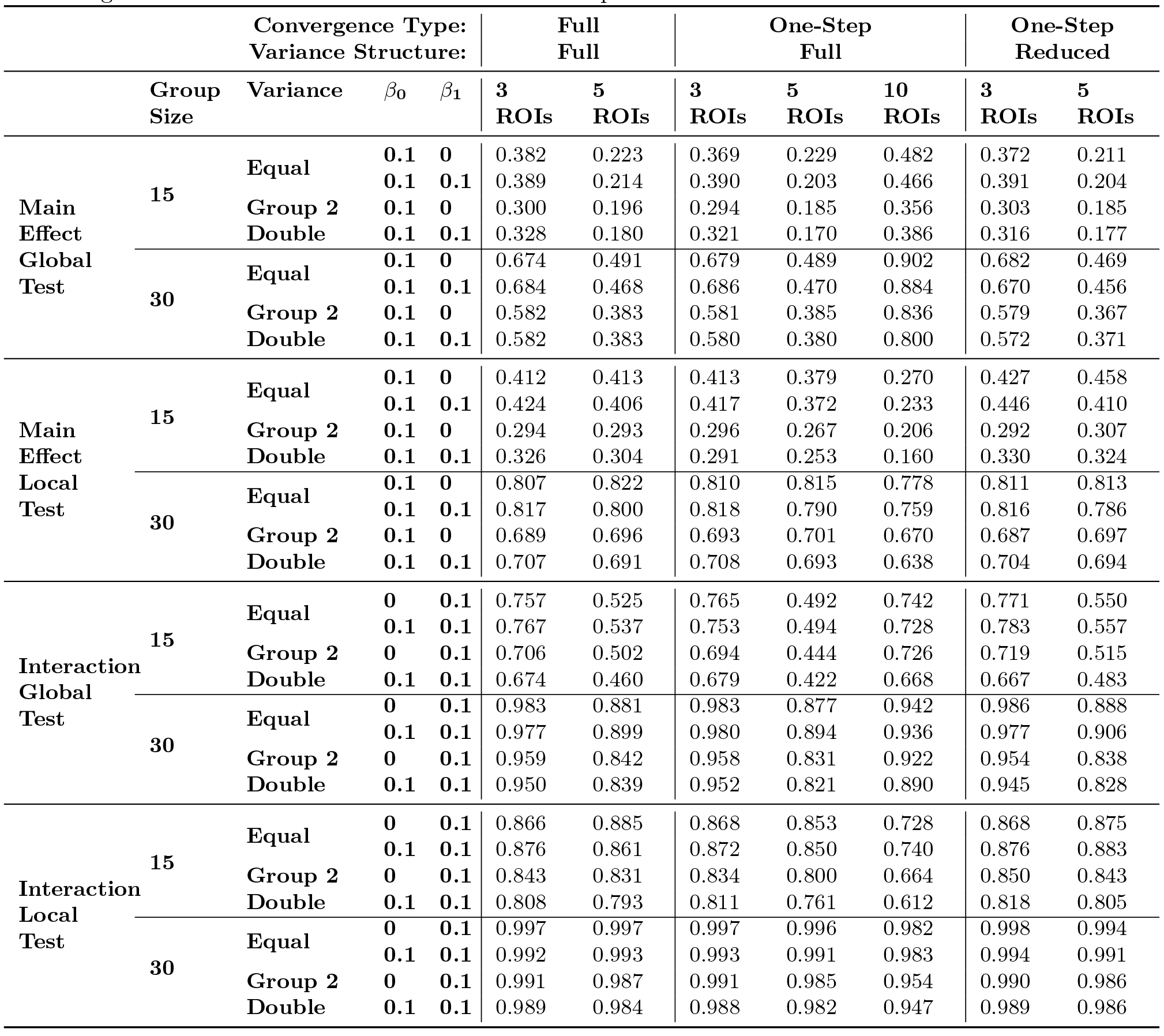
The power calculations for the simulation study. Power results for the main effect (group difference in intercepts) and interaction effect (group difference in longitudinal slopes) are reported both globally and locally. The global power results are averaged across all simulations. The local power results reported are unadjusted and averaged across all simulations and all non-null ROI pairs.

Overall, the full convergence results did not consistently improve on the one-step estimator results. With negligible gains in terms of power and Type I error, the additional computational resources used by the full-convergence model did not offer any practical advantage. A primary difference in the three models was the computational time. The full convergence model took, on average, over 2.5 times longer to run without seeing any boost in performance. For reference, using a 3.7 GHz Quad-Core Intel Xeon with 16GB ram, the average times to fit the one-step full variance model with 30 subjects per group for 3, 5, and 10 ROIs were 3.8 seconds, 54.9 seconds, and 87.2 minutes, respectively. These results show that the time increases quickly with the dimension of the model. The computational time is largely driven by the permutation procedure. Thus, if a larger number of permutations is desired for the testing procedure, then the computational time will see a corresponding increase.

## 4 ADNI Data Analysis

### 4.1 ADNI Preprocessing

Data used in the preparation of this article were obtained from the ADNI database (http://adni.loni.usc.edu). ADNI was launched in 2003 as a public-private partnership, led by Principal Investigator Michael W. Weiner, MD. The primary goal of ADNI has been to test whether serial magnetic resonance imaging, positron emission tomography, other biological markers, and clinical and neuropsychological assessment can be combined to measure the progression of MCI and early AD. For up-to-date information, see https://www.adni-info.org.

We preprocessed the ADNI data using both FSL (version 5.0.9, https://fsl.fmrib.ox.ac.uk/) and AFNI (version AFNI_17.0.15, https://afni.nimh.nih.gov/). The preprocessing steps were as follows. We 1) applied motion correction to the images using FSL’s mcflirt (rigid body transform; cost function normalized correlation; reference volume the middle volume) and then 2) normalized the images into the Montreal Neurological Institute space using FSL’s flirt (affine transform; cost function correlation ratio). We used FSL’s fast to 3) obtain a probabilistic segmentation of the brain to obtain white matter and cerebrospinal fluid (CSF) probabilistic maps, thresholded at 0.75. Using FSL’s fslmaths, we 4) spatially smoothed the volumes using a Gaussian kernel with FWHM=5 mm. We used AFNI’s 3dDetrend to 5) remove nuisance signals, namely the six motion parameters, white matter and CSF signals, and the global signal. Finally, 6) the linear trend was removed from each time series using linear regression and a 4th order Butterworth low-pass filter with a 0.1 Hertz cutoff was applied to each fMRI time series.

We used the Automated Anatomical Label (AAL) atlas to subdivide the brain into 116 anatomical regions [Tzourio-Mazoyer et al., 2002]. We define the ROI level time series for a given region as the average of the time series from each voxel (3D location) within that region of the brain. We then selected *P* =10 ROIs for analysis of the ADNI data based on previous literature which has shown differences in FC between AD and CN patients in the default mode network (DMN) and hippocampi [Supekar et al., 2008, Greicius et al., 2004, Sorg et al., 2007]. The ten regions we selected were the left and right hippocampus (HC), parahippocampus (PHC), posterior cingulate (PCC), precuneus (PQ), and prefrontal cortex (PFC). In all results that follow an l suffix for an ROI denotes the left side of the brain and an r suffix denotes the right side. Because a full brain analysis of all 116 regions is not currently feasible using our method, investigators should select a set of ROIs for their particular dataset and research question based on expert knowledge and literature review.

### 4.2 ADNI Analysis

Four models were fit to the ADNI data with differing assumptions. Model 1 is a one-step estimation model which assumes compound symmetry structure for Ψ_0_ and Ψ_1_ and unstructured Σ_*ij*_. Model 2, makes the same assumptions for Ψ_0_, Ψ_1_, and Σ_*ij*_ but uses the full convergence estimator. Model 3 is a one-step estimation model assuming scaled identity structures for Ψ_0_ and Ψ_1_ and a diagonal structure for Σ_*ij*_. Finally, Model 4 uses one-step estimation, assumes a diagonal structure for Ψ_0_, sets all elements of Ψ_1_ to 0, and assumes a diagonal structure for Σ_*ij*_. This final model is equivalent to a massive univariate approach which ignores the within-subject dependence. 5,000 permutations were run for all models. The intercept of each model represents the FC network of each group at age 65.

### 4.3 ADNI Results

Table 3 shows results from the global hypothesis tests and all local hypothesis tests that were significant before *p*-value adjustment for all four models. Neither the overall main effect or interaction term were found to significantly differ between groups in the global tests in any of the four models considered. The only ROI pair level difference that remained significant after *p*-value adjustment and correction in any of the models was the difference in the CN and AD group longitudinal slopes in the FC between the left HC and the right PCC in Models 1 and 2. These models conclude that the FC between HCl and PCCr declines at a significantly quicker rate in the AD population than in their CN counterparts. The estimated Model 1 group intercepts, group longitudinal trends, group differences in intercepts and longitudinal trends, and – log_10_ *p*-values after correction and adjustment from local hypothesis tests are presented graphically in Figure 3. Similar figures for Models 2, 3, and 4 can be found in Web Figures 1, 2, and 3, respectively, of the web-based supplementary materials. The four models fit to the ADNI data present slightly different results. The difference between Model 1 and Model 2 is very minimal. The nearly identical results provide further support of the conclusion from the simulation study that the full convergence and one-step estimator models lead to very similar estimates and inference. Some more pronounced differences in results arise when Model 1 is compared with Models 3 and 4. These differences make sense considering that Model 4 does not account for the within-subject dependence of the FC and thus appears to suffer from diminished power to detect group differences.

**Figure 3:**
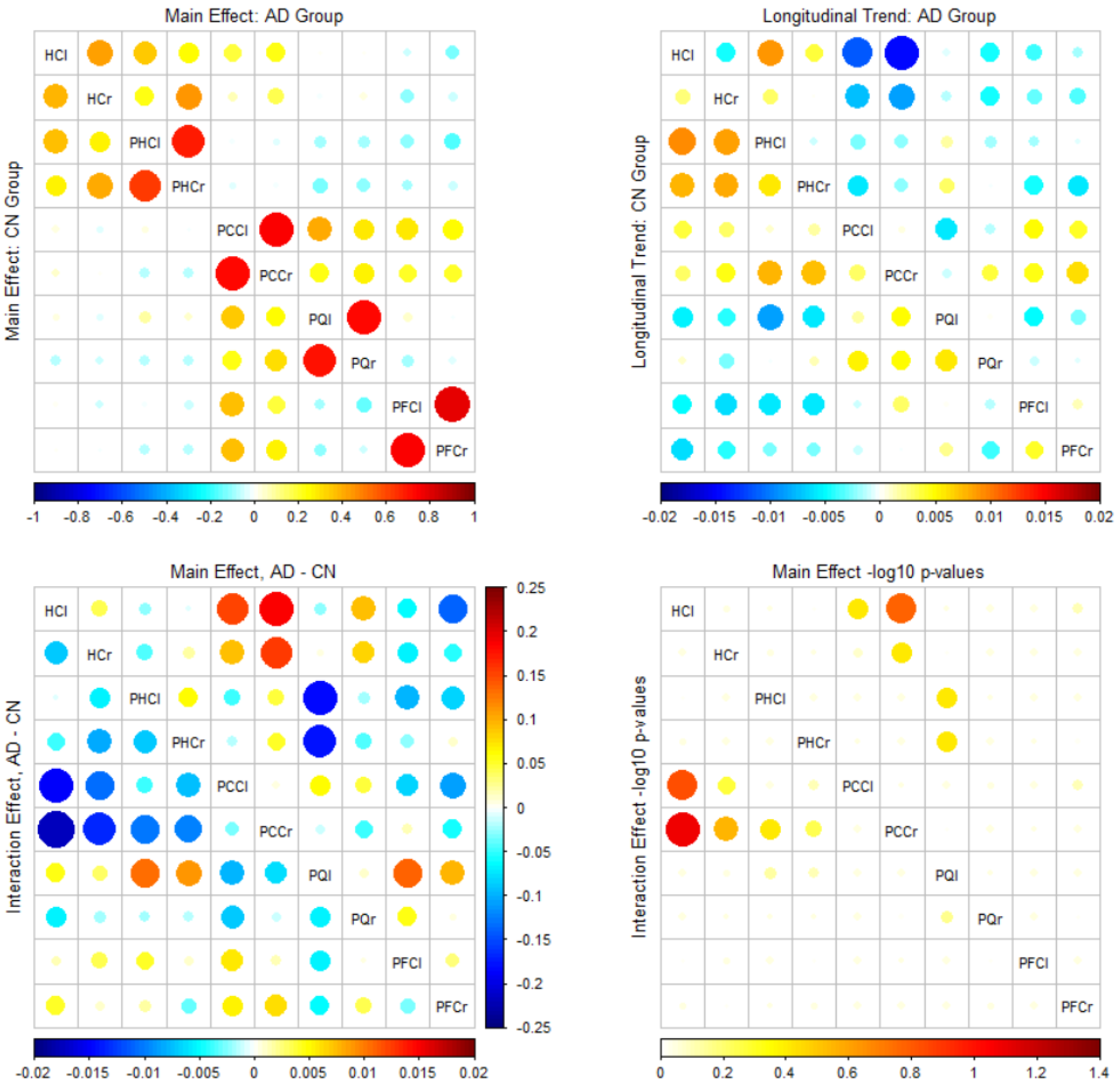
Model 1 results. Top left: A plot of the estimated intercept terms for the CN group (bottom left triangle) and AD group (top right triangle). Top right: A plot of the estimated slope terms for the CN group (bottom left triangle) and AD group (top right triangle). Bottom left: a plot of the group differences (AD estimates - CN estimates) for the estimated intercepts (top right triangle) and slopes (bottom left triangle). Bottom right: A plot of the — log_10_ corrected and adjusted *p*-values from all local hypothesis tests of group differences (AD estimates - CN estimates) for the estimated intercepts (top right triangle) and slopes (bottom left triangle).

The more interesting differences exist in comparing Models 1 and 3, but some common patterns run throughout both sets of results. In both models, many of the local hypotheses that were significant prior to the FDR correction appear between the HC/PHC and the PCC. These group differences strengthen the one local hypothesis that is significant after FDR correction from Models 1 and 2 which shows a significantly larger decrease in FC between the HCl and PCCr in the AD group than in the CN group. While the significant results disappear after FDR correction for Model 2, the fact that many other HC/PHC connections with the PCC see a similar pattern helps to indicate differing baseline and longitudinal trend effects in the FC of the two groups. This clustering of group differences can be seen in Figure 3 with the smallest *p*-values (red and orange circles) appearing between the HC/PHC and PCC. Wang et al. [2006], Sorg et al. [2007], and Greicius et al. [2004] all noted decreased FC between the HC and PCC in patients with AD in analyses of cross-sectional data. Similar results from Supekar et al. [2008] showed decreased clustering coefficients for the HC. Our analysis confirms these results with the addition of a longitudinal component to the analysis. Our results not only conclude that AD and CN patients have differing FC between the HC and PCC, as the previous works have shown, but we also more clearly describe the differences in baseline and longitudinal trend in FC between these two regions.

**Table 3:**
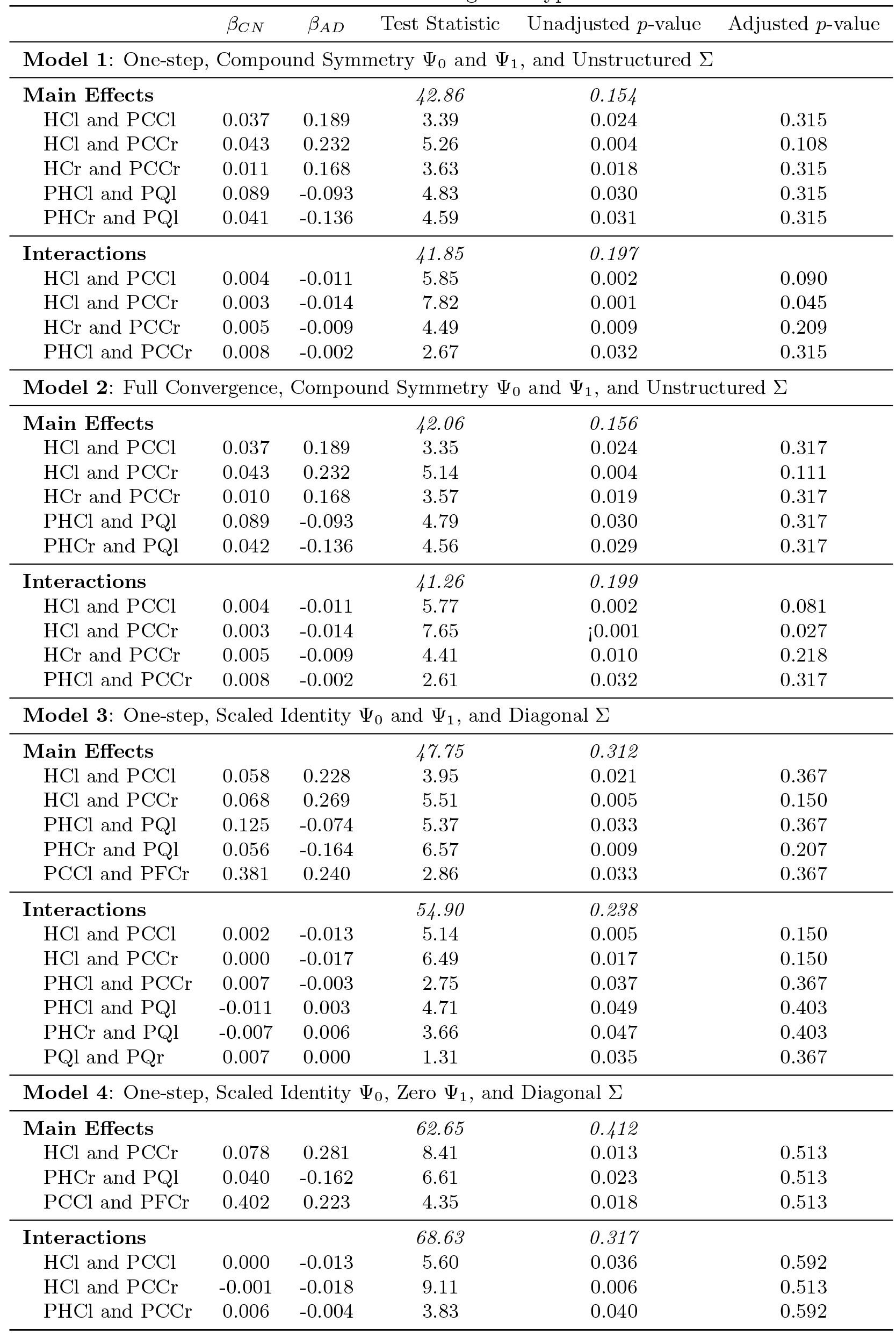
Hypothesis tests on the ADNI data. Global tests and all local tests with unadjusted *p*-values of < 0.05 are shown for Models 1-4. The numbers in italics are from the global hypothesis tests.

## 5 Discussion and Conclusions

We have introduced a novel variance components longitudinal model to estimate and draw inference on the group differences in FC using resting-state fMRI data. The model properly accounts for the correlation inherent in repeated measures data and the autocorrelation present in fMRI time series. A permutation testing procedure performs global and local two-sample testing in an efficient manner. The linear model framework and utilization of generalized least squares estimators offers great simplicity and a large number of natural extensions. This work fills a current gap in the literature by providing a general framework for estimation and hypothesis testing of longitudinal FC data.

As a practical example, we applied the method to resting-state fMRI data from the ADNI database. Our analysis found a faster decline in FC between the HCl and the PCCr in AD patients compared to the CN controls. This finding confirms the results of previous studies and helps solidify the central roles of the hippocampus and default mode network in AD.

Our familiar linear model framework allows for easy adoption and understanding of the model and its results. Additionally, the linear model framework offers many natural extensions. One could easily include terms for additional covariates such as scanner effect or gender. Different structures for the variance components could also be implemented to capture a wider range of possible correlation structures.

Our current method has the advantage of allowing for joint modeling of a complete FC network rather than taking a massive univariate approach. We see this joint modeling as a significant step forward, but complete brain analyses are still not yet feasible due to high computational demands of a model fit to many ROIs and the limited sample size of many fMRI studies. Here we have fit models to 10 ROI networks, but many brain atlases include more than 100 regions. In the future, some form of regularization could be introduced into the model to allow for analysis of an entire brain atlas. The ability to run a full brain FC network analysis would alleviate the issues arising from ROI selection.

The selection of the proper structure for the variance components deserves more attention. While a block compound symmetry structure for Ψ has a natural interpretation similar to that of a random intercept, there are certainly other viable structures. Choosing between structures is not a trivial task. One way to alleviate the model selection dilemma is to introduce a more robust sandwich type estimator of 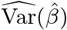, in which case incorrect specification of the variance would lead to valid inference with only a reduction in power.

## 6 Acknowledgments

Data used in preparation of this article were obtained from the ADNI database (adni.loni.usc.edu). As such, the investigators within the ADNI contributed to the design and implementation of ADNI and/or provided data but did not participate in analysis or writing of this report. A complete listing of ADNI investigators can be found at: http://adni.loni.usc.edu/wp-content/uploads/how_to_apply/ADNI_Acknowledgement_List.pdf.

